# A robust and scalable image analysis framework for phenotyping human thoracic aorta

**DOI:** 10.64898/2026.10.11.758402

**Authors:** Jiasong Chen, Linchen Qian, Ruonan Gong, Christina Sun, Thuy Pham, Caitlin Martin, Mohammad Zafar, John Elefteriades, Wei Sun, Liang Liang

**Affiliations:** Department of Computer Science, University of Miami, Coral Gables, FL, USA; Sutra Medical Inc., Lake Forest, CA, USA; Aortic Institute at Yale-New Haven Hospital, Yale University School of Medicine, New Haven, CT, USA

## Abstract

Large-scale phenotyping of the thoracic aorta requires a workflow in which every step, from segmentation to phenotype extraction, runs without manual intervention. We present a robust and scalable framework that converts contrast-enhanced CT into anatomical landmarks, a patient-specific quadrilateral mesh, a wall-stress field and regional phenotypes of the thoracic aorta. Four nnU-Net segmentation tasks delineate the aorta, its anatomical regions and two partitions of the aortic root. A surface–curve–point procedure derives the annulus, the sinotubular junction, the ostia of the supra-aortic branches, the cusp nadirs and the commissures from the interfaces between segmented regions. We fit a structured template of 24,882 quadrilateral elements to the segmented aortic surface by staged, differentiable mesh deformation and compute the transmural mean wall stress with PyTorch-FEA. The structured mesh provides node-to-node correspondence across patients, anchored to the anatomy by the landmarks, from which the framework extracts regional morphological and biomechanical phenotypes.

## Introduction

Maximal aortic diameter is the primary imaging criterion for surveillance and elective repair of thoracic aortic aneurysm, with guidelines combining diameter thresholds with growth rate and body-size-indexed dimensions^1^. Diameter is simple to measure and to report, but it reduces a three-dimensional structure to a single number. It does not describe regional size, curvature or tortuosity, and acute type A dissections can occur at diameters below the threshold for elective repair^2^.

Morphological and biomechanical measures can complement diameter. Morphological measures quantify the three-dimensional geometric shape of each aortic region. Biomechanical measures quantify the mechanical state of the wall under blood pressure, which finite element analysis (FEA) can compute from patient-specific geometry^3,4^. Direct measurement is not an alternative. The mechanical properties and the strength of the aortic wall are measured ex vivo on tissue excised during open surgery^5^, so they exist for few patients and are not available before the decision to operate. Image-based simulation is therefore a noninvasive route to wall mechanics, and it can be applied to every patient who has a CT scan.

An image-to-phenotype workflow has several steps, and each step must be automated to enable large-scale analysis. The first step is segmentation. Self-configuring networks such as nnU-Net segment the aorta in contrast-enhanced CT automatically^6^. The second step is anatomical landmark localization. Regional analysis and mesh construction need landmarks: the aortic annulus, the sinotubular junction (STJ), the cusp nadirs and commissures of the aortic root, and the ostia of the supra-aortic branches. Manually placing these landmarks on every scan does not scale to large populations. The third step is mesh generation, the fourth is mechanical analysis, and the fifth is phenotype extraction. Open-source tools cover vascular geometry processing^7^ and blood-flow simulation^8^, whereas a wall-stress analysis also needs a structured mesh of the wall and a solid-mechanics solver.

Here we present an automated framework that converts contrast-enhanced CT into anatomical landmarks, a patient-specific quadrilateral mesh, a wall-stress field and regional phenotypes of the thoracic aorta, from the aortic annulus to the level of the celiac trunk. The framework has five steps: segmentation with nnU-Net^6^, landmark extraction from the segmentation, simulation-ready remeshing by fitting a structured template, wall-stress computation, and extraction of regional morphological and biomechanical phenotypes. The landmark extraction requires no manual annotation at inference: it derives every curve and point from the anatomical interfaces between segmented regions. All patients with the same arch variant share one template topology. We describe the method and report preliminary results: landmark extraction, remeshing and wall stress in two example patients, and a summary of the phenotypes in the study population.

## Methods

### Data and reference annotation

We selected 660 contrast-enhanced CT scans from the public INSPECT dataset^9^. We excluded scans without contrast enhancement, scans that did not cover the thoracic aorta and scans of poor image quality. The selection balanced sex, aortic diameter range and arch variant: a three-branch arch, or a two-branch (bovine) arch, in which the brachiocephalic trunk (BT) and the left common carotid artery (LCCA) share one ostium. All selected patients had tricuspid aortic valves, and we excluded four-branch arches. The scans were split into 526 training cases and 134 test cases, with the diameter ranges in equal proportions in the two sets.

Three annotators established the reference annotation by consensus in 3D Slicer^10^. The annotation comprised segmentation labels, root landmarks and anatomical boundary curves. The whole-aorta label was divided into seven regions. The left ventricular outflow tract (LVOT) region was a 5 mm extension on the ventricular side of the annulus plane. The aortic root extended from the annulus plane to the STJ, excluding the coronary arteries. The thoracic aorta comprised the ascending aorta, the arch and the descending aorta. The proximal abdominal region ended at a plane about 5 mm distal to the origin of the celiac trunk. The other three regions were the arch branch segments, about 5–10 mm long, of the BT, the LCCA and the left subclavian artery (LSA). The root landmarks were the three cusp nadirs, defined as the most basal points of the leaflet attachment lines, and the three commissures. We applied two partitions to the root. The sinus partition divided the root into three sinus sectors, one for each of the left coronary, right coronary and non-coronary sinuses of Valsalva, with boundaries through the commissures. The commissural partition divided it into three commissural sectors, each centered on one commissure, with boundaries through the cusp nadirs. Each commissural sector contains one interleaflet triangle and the adjacent halves of the two neighboring sinuses.

### Segmentation

We trained one nnU-Net^6^ for each of four segmentation tasks, in the 3D full-resolution configuration: the whole aorta, the seven regions, the sinus partition of the root and the commissural partition of the root. For the first two tasks, images were resampled to 1.0 mm isotropic spacing. For the two root-partition tasks, we cropped the image to 1.2 times the bounding box of the root and resampled the crop to 256 × 256 × 256 voxels. Intensities were clipped at the 0.5th and 99.5th percentiles and scaled to the range 0–1. We trained each network with five-fold cross-validation within the training set (420–421 training and 105–106 validation cases per fold) and used the model with the best validation performance for inference. At inference, the whole-aorta and seven-region networks ran on the resampled image. The predicted root region defined the cropping volume on which the two root-partition networks were applied. We kept the crop transformation, so that structures found in the crop could be mapped back to original image coordinates, and removed small and disconnected components from every predicted mask.

### Landmark extraction from the segmentation

We extracted the anatomical landmarks directly from the segmentation masks in three steps, from surfaces to curves to points (Fig. 1a). No step uses image intensities or manual input.

**Fig 1.**
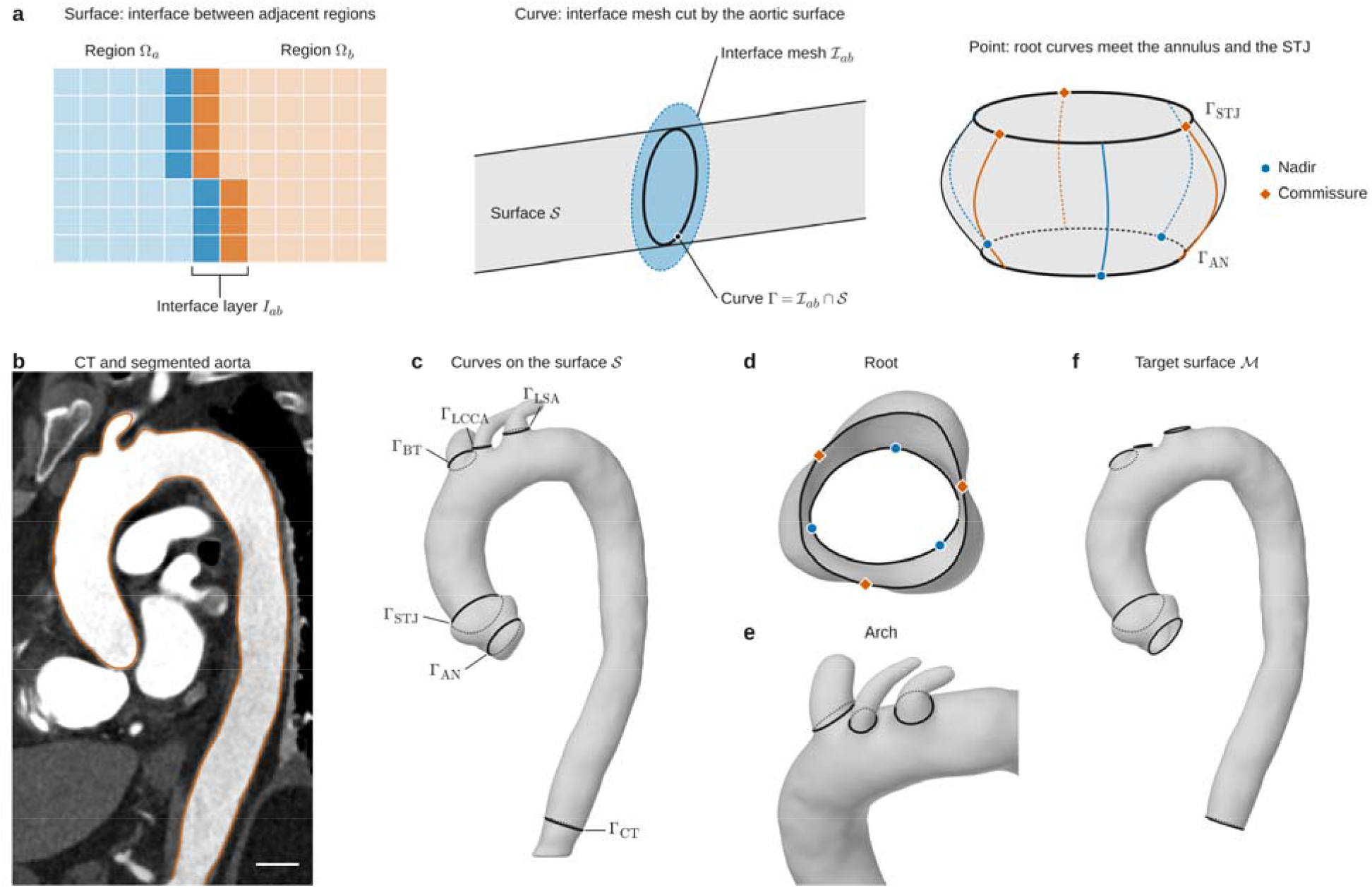
Landmark extraction from the segmentation. **a**, Schematic of the surface–curve–point procedure. Left, the voxels of two adjacent label regions that touch the other region form the interface layer. Middle, the interface layer is converted to a surface mesh and enlarged by 10% about its centroid (dashed outline); its intersection with the whole-aorta surface *S* is an anatomical curve. Right, in the root, the curves obtained from the two root partitions meet the annulus curve at the three cusp nadirs and the sinotubular junction (STJ) curve at the three commissures. **b**, CT (display range −200 to 600 HU) with the outline of the segmented aorta (orange). **c**, Surface *S* with the six anatomical curves. **d**, Root between the annulus curve and the STJ curve, viewed along the root axis from the STJ side, with the nadirs (blue) and commissures (orange). **e**, Arch with the three branch curves. **f**, Target surface ℳ, obtained by cutting *S* along the annulus, celiac trunk level and branch curves. AN, annulus; BT, brachiocephalic trunk; CT (subscript), celiac trunk level; LCCA, left common carotid artery; LSA, left subclavian artery.

#### Surfaces

We converted the whole-aorta mask into a triangular surface mesh *S* in physical coordinates with marching cubes^11^. For each pair of adjacent regions with masks *Ω*_*a*_ and *Ω*_*b*_, the interface layer *I*_*ab*_ consists of the voxels of each region that touch the other. Marching cubes converted *I*_*ab*_ into an interface mesh *ℐ*_*ab*_, which we enlarged by a factor of 1.1 about its centroid so that it extends beyond the aortic wall.

#### Curves

The intersection of an interface mesh with *S* is a closed curve on the aortic surface. The interfaces of the seven-region segmentation give the annulus *Γ*_AN_(LVOT and root), the STJ curve *Γ*_STJ_(root and thoracic aorta), the celiac trunk level curve *Γ* _CT_ (thoracic aorta and proximal abdominal region) and one branch curve around each ostium, *Γ* _BT_, *Γ*_LCCA_ and *Γ*_LSA_ (thoracic aorta and the branch; *Γ*_LCCA_ is absent in a two-branch arch). A branch interface can lie oblique to the branch axis. We therefore rotated each branch With *S*, so that the branch curve approximates a cross section perpendicular to the branch. The interfaces interface mesh about its centroid to the orientation that minimizes the area enclosed by its intersection of the two root partitions give six further curves on the root surface, computed in the cropped root frame and mapped back to original physical coordinates. The interfaces between the three commissural sectors give three curves that pass through the cusp nadirs (nadir curves), and the interfaces between the three sinus sectors give three curves that pass through the commissures (commissure curves).

#### Points and target surface

All curves lie on *S*. Each nadir point is therefore the intersection of a nadir curve with *Γ*_A N_, and each commissure point is the intersection of a commissure curve with *Γ*_STJ_. We resampled *Γ*_AN_ and *Γ*_STJ_ to 78 ordered points each, with the three nadirs at points 0, 26 and 52 and the three commissures at points 13, 39 and 65, which matches the node numbering of the template below. Cutting *S* along *Γ*_AN_, *Γ*_CT_ and the branch curves and keeping the part between *Γ*_AN_ and *Γ*_CT_ gave the target surface *ℳ* for remeshing. We evaluated the root landmarks against the reference annotation by the distance between the automated and reference points, for the nadirs and for the commissures.

### Template

For each patient, aortic centerlines were extracted using VMTK^7^ and used to guide the construction of patient-specific tubular templates. The tube has 320 cross sections of 78 nodes each (24,960 nodes and 24,882 quadrilateral elements). Resampling the centerline at uniform arc length between landmarks placed the annulus, the STJ, the BT branch and the celiac trunk level on cross sections 0, 20, 80 and 319. In a three-branch arch, the LCCA and the LSA lie on cross sections 100 and 120; in a two-branch arch, the LSA lies on cross section 100 (Fig. 2a). The landmarks also set the circumferential orientation. Nodes 0, 26 and 52 of cross section 0 correspond to the three nadirs, and nodes 13, 39 and 65 of cross section 20 to the three commissures. Between these cross sections, a rotation-minimizing frame^12^ carries the orientation along the centerline. At each branch, we opened the tube within the 12 × 12-element patch centered on node 39 of the branch cross section. We removed a central square the size of 3 × 3 elements to form the ostium, placed 48 nodes on its border and connected them to the patch border by three rings of 48 elements (Fig. 2b). Each opening leaves the element count unchanged, so the template *ℳ*_o_ has 24,882 elements and 25,029 nodes (three openings) or 25,006 nodes (two openings).

**Fig 2.**
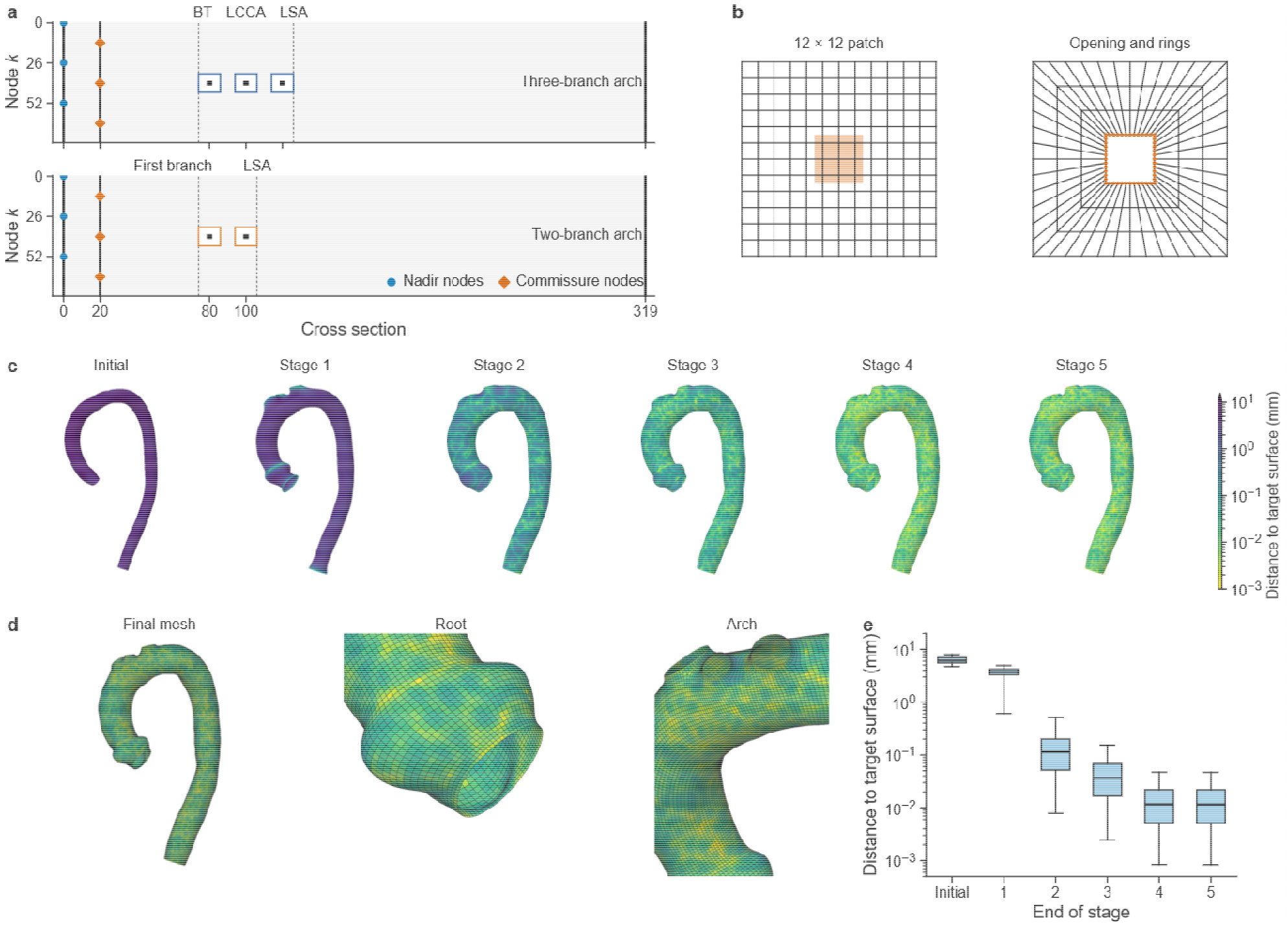
Simulation-ready remeshing. **a**, Template layout for the two arch variants: cross section by circumferential node *k*. Thick lines, cross sections constrained to the annulus (0), sinotubular junction (20) and celiac trunk level curve (319); squares, 12 × 12-element ostium patches with the opening in black; dashed lines, aortic arch region cross sections. BT, brachiocephalic trunk; LCCA, left common carotid artery; LSA, left subclavian artery. **b**, Opening construction: the central square of the patch (orange) is removed, and three rings of 48 elements connect the 48 border nodes of the opening (orange) to the patch border. **c**, Mesh with openings of patient 1 (three-branch arch) at the start of fitting and at the end of each stage, colored by node distance to the target surface (logarithmic scale); left anterior oblique view. **d**, Final mesh of patient 1. **e**, Node distance to the target surface for the 25,029 nodes at the end of each stage.

### Template fitting

We fitted *ℳ*_0_ to *ℳ* by optimizing nodal displacements with the element connectivity fixed. At iteration *k*, the node positions are **X**^(*k*)^=**X**_0_+**U**^(*k*)^ where **X**_0_ holds the initial nodal coordinates and **U**^(0)^= **0**. Large displacements under weak regularization can distort, fold or invert elements. We therefore moved the constrained nodes to their curves in small increments. We constrained six node sets to the anatomical curves: the 78 nodes of cross sections 0, 20 and 319 to *Γ*_AN_, *Γ*_STJ_ and *Γ*_CT_, and the 48 border nodes of each ostium to its branch curve. Let *ℬ* be the set of constrained nodes, **x**_*Γ,v*_ the target point of node *v* ∈ ℬ on its curve, and ℱ the set of free nodes. The template fitting ran for *K* iterations in *S* stages, which differed in the loss weights and the learning rate. Stage s ended at iteration *k*_S_, with *k*_0_=0 and *k*_S_ =*K*. In stage s, we minimized the loss ℒ ^(s)^ over the free-node displacements **U** _ℱ_ with prescribed displacements at the constrained nodes:

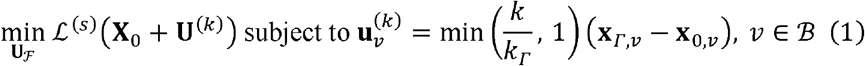

For *k* _s−1_ < *k* ≤ *k*_s_. **x**_0,*v*_ and 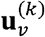 are the initial position and the displacement of node *v* ∈ ℬ and *k* _*Γ*_ *≤K* is the number of iterations over which the constrained nodes approach their curves. The constrained nodes thus reach their target points in *k*_*Γ*_ equal increments and stay there for *k> k*_*Γ*._ In this study, we used *S*=5 stages and *K*=14,000 iterations, and the boundary approach spanned the first stage(*k*_*Γ*_=*k*_1_). *ℒ*^(s)^is the mesh loss with the weights of stage *s:*

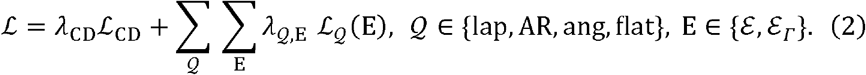

ℒ_CD_ is a Chamfer distance between the template nodes and the target surface ℳ, and it draws the template mesh onto ℳ. The quality terms act on all elements ℰ and, with separate weights, on the elements ℰ _*Γ*_ next to the constrained boundaries. The Laplacian termℒ_lap_ penalizes the squared distance of each node from the mean position of its edge-connected neighbors, which keeps the surface smooth. The aspect ratio termℒ_AR_ penalizes unequal edge lengths within an element, the angle term ℒ_ang_ penalizes the deviation of each interior angle from its target (90° at regular nodes), and the flatness term ℒ_flat_ penalizes non-planar elements. The Laplacian weight was high in stages 1–3 and reduced by a factor of ten in stages 4 and 5, analogous to the progressive reduction of stiffness in non-rigid registration^13^.Stronger regularization favored smooth deformation in the early stages, whereas weaker regularization allowed finer adaptation to local anatomy. Adamax^14^ updated **U**_ℱ_ at a learning rate of 0.05, reduced to 0.01 from stage 4, with gradients from automatic differentiation in PyTorch^15^ . We used one set of weights and one schedule for all patients. As a simpler variant for FEA, we also fitted the tube without openings to ℳ with each ostium sealed (*K* = 13,000).

### Wall-stress computation

We computed wall stress with PyTorch-FEA, our open-source finite element solver for aortic mechanics^4^. CT does not provide the material properties of the patient’s aortic wall. For a thin-walled vessel under pressure, however, force equilibrium mainly sets the stress averaged across the wall thickness, and this mean stress depends only weakly on the material, a property known as statical determinacy^3,16,17^. Its simplest example is Laplace’s law for a thin-walled cylinder, in which the circumferential stress equals the pressure times the radius divided by the wall thickness. We modeled the wall as a stiff neo-Hookean material, so that the computed stress depends on geometry and pressure and not on the unknown tissue properties^4^. From each fitted mesh, we built a solid wall of uniform 2 mm thickness with four layers of eight-node hexahedral elements, taking the fitted surface as the mid-wall surface. We extended both ends of the wall by 1 mm and fixed all nodes at the two ends. A uniform static pressure *p* = 16 kPa (120 mmHg) acted on the inner (luminal) surface of the wall in every model, so that under blood pressure, we treated the solid model as the pressurized state. Let x denote the nodal positions stress differences between patients reflect differences in aortic geometry. Because CT shows the aorta of the solid wall in the pressurized state and *X*_*zp*_ the nodal positions of the zero-pressure geometry. We solved one inverse analysis for the unknown *X*_*zp*_ such that the wall is in equilibrium in the pressurized state:

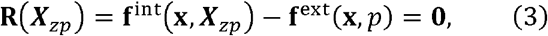

at all nodes that are not fixed, where **R** is the residual force, **f**^int^ are the internal nodal forces arising from the Cauchy stress *σ*, and **f**^ext^ are the external nodal forces of the pressure *p*. The stress *σ*follows from the deformation gradient *F* = ∂x/ ∂*X*_zp_ through the neo-Hookean law. PyTorch-FEA solved equation (3) by energy minimization with L-BFGS^18^ and adaptive load increments^4^. Once *x*_*zp*_ is known, the stress field *σ* in the pressurized state follows directly from ***F***. For each element *e* of the fitted surface, the transmural mean principal stress is the average over the four layers of the wall:

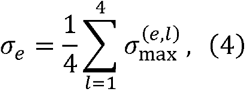

where 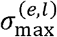 is the principal Cauchy stress of largest magnitude in the hexahedral element of layer *l* at *e*. We discarded the stress on the extensions and assigned each node the mean stress value of the elements that share it. Below, wall stress refers to *σ* at the nodes.

### Regional phenotypes

The last step characterizes each aortic region by morphological and biomechanical phenotypes. It uses the fitted mesh without openings and its *σ* at 16 kPa. Because all fitted meshes share node correspondence, cross-section indices define the regions once for all patients: the root (cross sections 0–20), the ascending aorta (20–70), the arch (70–126 in a three-branch arch, or 70–108 in a two-branch arch) and the descending aorta (to 319). Along the aorta, we cut the fitted surface with planes perpendicular to the centerline and computed the area, the perimeter and the diameter. Each region is further described by its centerline length, tortuosity index (the centerline length divided by the straight distance between its ends, minus one, in percent), lumen volume, wall area, peak *ℳ*, among other phenotypes.

## Results

### Segmentation and landmarks

We report the preliminary results for two example patients from the test set. patient 1 (three-branch arch) and patient 2 (two-branch arch). In both patients, the landmark points came from the automatic extraction without manual correction. Over the two patients, the mean distance between the automated and reference points was 0.31 mm for the six cusp nadirs and 0.44 mm for the six commissures.

Figure 1b–f shows the landmark extraction in patient 1, who has a three-branch arch. The six anatomical curves lay on the whole-aorta surface (largest distance to the nearest surface vertex 0.18 mm). The annulus and STJ curves enclosed the root, the three nadirs lay on the annulus curve and the three commissures on the STJ curve (Fig. 1d). Each branch curve encircled the base of one supra-aortic branch (Fig. 1e), and cutting along the curves removed the branches and left one opening per ostium in the target surface (Fig. 1f).

### Remeshing and wall stress

In patient 1, the initial template lay a median of 6.1 mm from . The median distance fell to 3.7 mm at the end of stage 1, 0.11 mm at the end of stage 2, 0.037 mm at the end of stage 3 and 0.012 mm at the end of stages 4 and 5 (Fig. 2c,e). The final meshes with openings lay a mean of 0.016 mm (patient 1) and 0.013 mm (patient 2) from, with maxima of 0.26 mm and 0.32 mm (Fig. 2d; Table 1). For comparison, the image spacing was 1.0 mm. The largest deviation of an interior angle from its target had a median over elements of 2.3° and 1.6° and a 95th percentile of 10.9° and 7.6°. The largest deviations occurred between adjacent branch openings. In patient 1, the angle deviation exceeded 30° in 82 of 24,882 elements (maximum 52.6°), all between the BT and LCCA openings. In patient 2, it exceeded 30° in four elements (maximum 35.9°).

**Table 1.** Remeshing, wall stress and example phenotypes in the two example patients.

|  | Patient 1 | Patient 2 |
| --- | --- | --- |
| Arch variant | Three-branch | Two-branch (bovine) |
| Nodes; elements (mesh with openings) | 25,029; 24,882 | 25,006; 24,882 |
| Distance to $\mathcal{M}$ (mm): mean; median; 95th percentile; maximum | 0.016; 0.012; 0.047; 0.26 | 0.013; 0.009; 0.037; 0.32 |
| Angle deviation ($^{\circ}$): median; 95th percentile; maximum | 2.3; 10.9; 52.6 | 1.6; 7.6; 35.9 |
| $\sigma$, root (kPa) | 94.9 (68.6–119.6) | 95.0 (73.5–112.1) |
| $\sigma$, ascending aorta (kPa) | 118.5 (105.4–135.5) | 111.2 (100.9–124.6) |
| $\sigma$, arch (kPa) | 119.0 (108.0–129.8) | 102.6 (89.8–110.6) |
| $\sigma$, descending aorta (kPa) | 95.2 (88.3–105.2) | 77.9 (75.6–82.0) |
| $\sigma$, 99th percentile, whole aorta (kPa) | 159.2 | 142.7 |
| Centerline length (mm); tortuosity index (%) | 365.5; 246 | 343.7; 293 |
| Wall area (cm<sup>2</sup>); lumen volume (ml) | 314.8; 211.6 | 265.7; 162.7 |
| Maximum diameter (mm) | 37.1 | 34.1 |

Distances are point-to-surface distances from the nodes of the mesh with openings to the target surface $\mathcal{M}$ . Angle deviation is the largest deviation of an interior angle of an element from its target, on the mesh with openings. $\sigma$ values are the median (IQR) over the nodes of each region on the mesh without openings at 16 kPa. Centerline length, tortuosity index, wall area and lumen volume refer to the fitted mesh from the annulus to the celiac trunk level;

All four wall-stress simulations (two meshes in each patient) converged without retries. The zero-pressure and image-derived geometries differed by at most 0.10 mm at any node. The stiff wall therefore deformed negligibly under pressure, and the computed stress refers to the geometry seen in the image. On the mesh without openings, the median *σ* was 101.8 kPa (interquartile range (IQR) 90.9–118.6 kPa) in patient 1 and 81.2 kPa (76.4–98.9 kPa) in patient 2 (Fig. 3a– d). In both patients, the regional median of *σ* was higher in the ascending aorta and arch than in the root and descending aorta (Table 1; Fig. 3f,g). In the ascending aorta, arch and descending aorta, the ratio of the cross-section median of *σ* to the Laplace estimate *pr*/*t* had regional medians of 0.91–0.97, where *r* is the mean radius of the fitted cross section and *t* = 2 mm (Fig. 3h). A ratio slightly below 1 is expected, because the pressure acts on the luminal surface, 1 mm inside the fitted surface, which gives (*r* − m m)/= 0.91–0.94 for these radii. The computed stress was therefore consistent with Laplace’s law in the tubular segments. The ratio was lower in the root (0.80 and 0.83), where the cloverleaf cross section of the sinuses is not circular. The meshes with and without openings gave nearly the same stress (median absolute difference 0.2 kPa in patient 1 and 0.1 kPa in patient 2; 95th percentile 3.3 kPa and 1.5 kPa).

**Fig 3.**
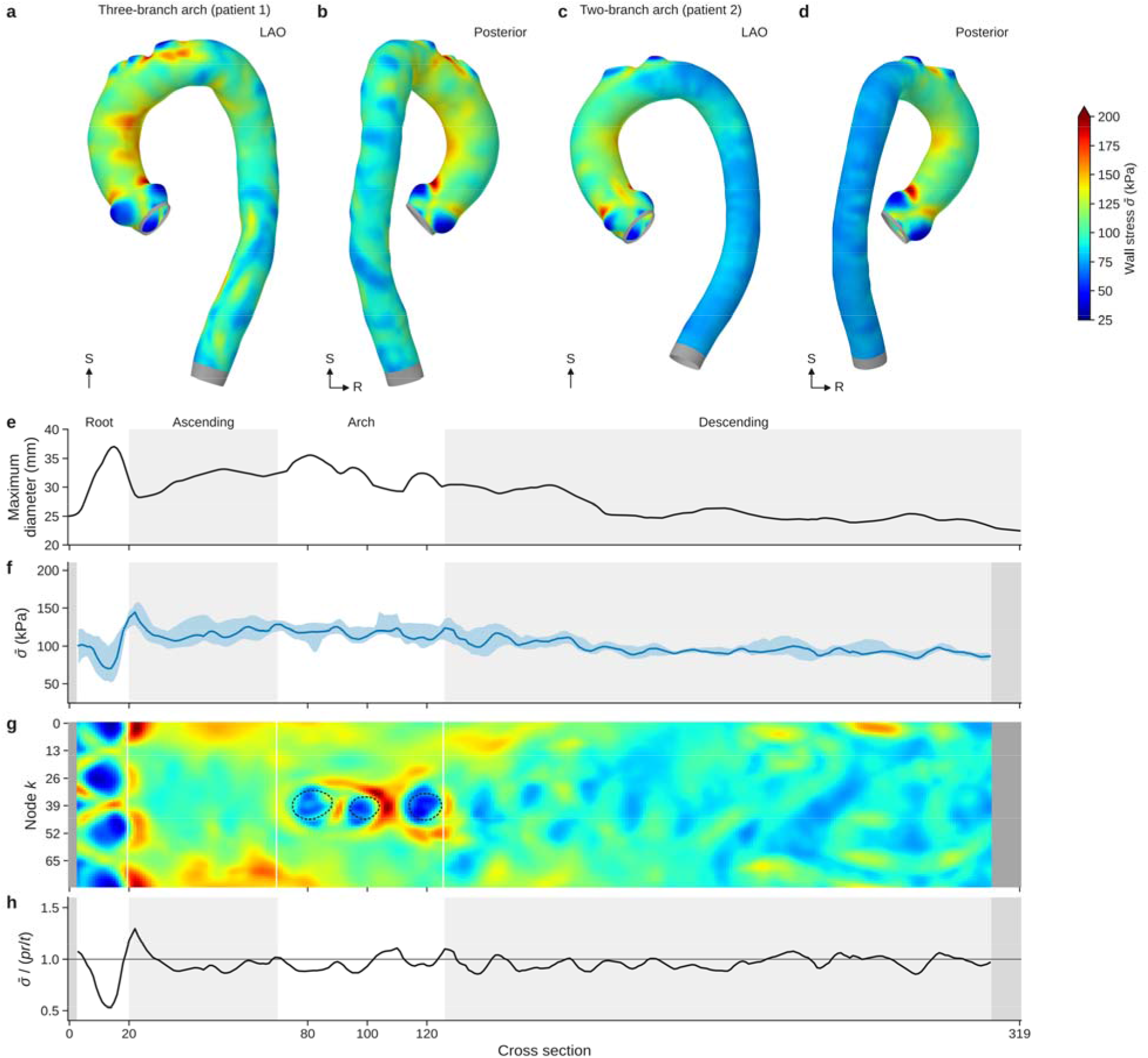
Wall stress computed on the fitted mesh follows Laplace’s law in straight segments and departs from it where the geometry changes. **a**–**d**, Transmural mean principal stress *σ* at 16 kPa; **a**,**c** left anterior oblique (LAO) view; **b**,**d** posterior view. **e**, Maximum diameter along the aorta of patient 1. **f**, *σ* along the aorta of patient 1. **g**, *σ* colormap of patient 1; dashed outlines, borders of the branch openings. **h**, Ratio of the cross-section median *σ* to the Laplace estimate *pr*/*t*; gray line, ratio 1.

### Phenotypes

In the two example patients, the fitted meshes gave centerline lengths of 365.5 mm and 343.7 mm from the annulus to the celiac trunk level, lumen volumes of 211.6 ml and 162.7 ml and maximum diameters of 37.1 mm and 34.1 mm (Table 1; Fig. 3e). For the study population, we extracted the phenotypes of 631 of the 660 patients and the framework generated 191 morphological and 57 biomechanical phenotypes per patient.

## Discussion

Without manual intervention at inference, the framework converted contrast-enhanced CT into anatomical landmarks, a quadrilateral mesh whose topology is fixed for each arch variant, and a wall-stress field. The framework treats landmark localization as the segmentation of anatomical regions. Every curve is derived from the interface between two segmented regions, and every root landmark is the intersection of two such curves. The landmark accuracy therefore depends only on the segmentation, and the framework needs no separate landmark detection network. This makes the landmark localization explainable and traceable: each landmark follows by geometric intersection from the label boundaries that define it.

In the two example patients, the mean errors of the nadirs and commissures were 0.31 mm and 0.44 mm, below the 1.0 mm image spacing. The fitted meshes followed the target surface to well within the image spacing, and the stress analyses converged without retries. Away from the root, the computed stress stayed close to the Laplace estimate, as expected for a thin-walled, nearly cylindrical vessel.

The shared template enables the meshes node-to-node correspondence, and the landmarks anchor this correspondence to the anatomy. In all patients, the same template cross sections lie at the annulus, the STJ and the celiac trunk level, and the same nodes lie at the nadirs and commissures. Shape and stress can therefore be compared node by node across patients and across longitudinal scans of one patient. Anatomical regions are fixed sets of node indices, which is the basis of the regional phenotypes. This framework could be extended to hemodynamic analysis, with the branch ostial openings serving as outlet boundaries for computational fluid dynamics simulations.

## Notes

### Competing Interest Statement

The authors have declared no competing interest.

